# DeepNose: Using artificial neural networks to represent the space of odorants

**DOI:** 10.1101/464735

**Authors:** Ngoc Tran, Daniel Kepple, Sergey A. Shuvaev, Alexei A. Koulakov

## Abstract

The olfactory system employs an ensemble of odorant receptors (ORs) to sense odorants and to derive olfactory percepts. We trained artificial neural networks to represent the chemical space of odorants and used that representation to predict human olfactory percepts. We hypothesized that ORs may be considered 3D spatial filters that extract molecular features and can be trained using conventional machine learning methods. First, we trained an autoencoder, called DeepNose, to deduce a low-dimensional representation of odorant molecules which were represented by their 3D spatial structure. Next, we tested the ability of DeepNose features in predicting physical properties and odorant percepts based on 3D molecular structure alone. We found that despite the lack of human expertise, DeepNose features led to perceptual predictions of comparable accuracy to molecular descriptors often used in computational chemistry. We propose that DeepNose network can extract *de novo* chemical features predictive of various bioactivities and can help understand the factors influencing the composition of ORs ensemble.

## Introduction

The olfactory system has evolved to sense and recognize a large number of chemicals present in the environment. Odorant receptors (ORs) act as molecular sensors that bind odorants and convey information about their structure to higher brain regions. ORs are activated by molecules in a combinatorial manner: one receptor binds to many molecules, and one molecule can interact with many receptors^1–4^. The number of ORs varies amongst species: from 350 in humans to about 1000 in most other mammals^5^. Here, we study the factors influencing the OR ensemble composition.

In this study, we use multilayer (deep) convolutional neural networks (CNNs)^6^ to study the space of molecules and use derived representations to predict human perceptual responses. CNNs are multilayer neural networks that use ensembles of spatial filters of increasing complexity to extract features relevant to representing objects present in the input^7^. Networks based on CNNs are the best performing method for image recognition, many of which perform better at classifying images than humans^8,9^. The performance of CNNs in 2D image recognition tasks suggests that they are well-suited for learning other types of spatial data, such as the 3D molecular structures. Deep networks have indeed been applied to classify molecules, including the predictions of pharmacokinetics properties, toxicity, and protein-ligand interaction^10–12^. CNNs were also used to represent the SMILES string of molecules^13^. In this study, we derive CNNs that can recognize 3D molecular structures and predict olfactory percepts.

Our main hypothesis is that OR ensemble forms a set of 3D filters that interact with molecular structures in real space. In this regard, molecules play the role of 3D images, and ORs can be viewed as analog of receptive fields, such as edge detectors, of the visual system^14^. Both visual receptive fields and the OR ensemble have been shaped by evolution to sense relevant features present in the stimuli. We thus hypothesize that *in silico* OR ensemble can be optimized using conventional machine learning methods, such as back-propagation^15^, to accurately represent the space of molecules. Using this strategy, machine learning may help us understand the factors influencing OR evolution and construct a model for OR-to-ligand binding that is not so remote from reality.

One of the biggest limitations in building a robust predictive computational model of olfactory processing is the lack of specific data on OR binding affinities and perceptual odorant descriptors. Recent efforts have been successful in generating data for hundreds of odorants^16^, while modern machine learning techniques require thousands to millions of data points to be successful^6^. To address this problem, we train a stacked deep autoencoder^17^, which we named DeepNose, to replicate 3D molecular structures received as inputs after the data is substantially compressed in the middle of the network. Training the autoencoder relies on unsupervised learning, and, as such, can be implemented using large scale unlabeled datasets containing 3D molecular structures.

We found that the autoencoder can be trained to fully and accurately replicate input molecules. Therefore, DeepNose can be viewed not only as a functional analog of the ORs ensemble, but also as a data-driven and reversible feature extractor. We further tested the biological relevance of the extracted features by predicting various biological activities. The resulting accuracies were comparable to results obtained with E-Dragon chemical descriptors, a state-of-the-art chemical feature extraction software widely used in odorant percept prediction^16,18^.

## Results

Molecules in the environment can be viewed as 3D objects that are sensed and recognized by the olfactory system. Similarly to visual images, each molecule can be represented by separate ‘color’ channels that contain information about the spatial distribution of six elements: C, H, O, N, S, and Cl. This forth discrete dimension will be called here ‘CHONSCl’, similarly to RGB channels used in images. In contrast to the visual system, olfactory objects can be presented to the sensory system in multiple copies simultaneously, differing in orientation and conformations. We therefore represent each input molecule by a 5D object, including with three spatial dimensions, one ‘color’ CHONSCl, and one orientation dimension (Figure 1).

**Figure 1.**
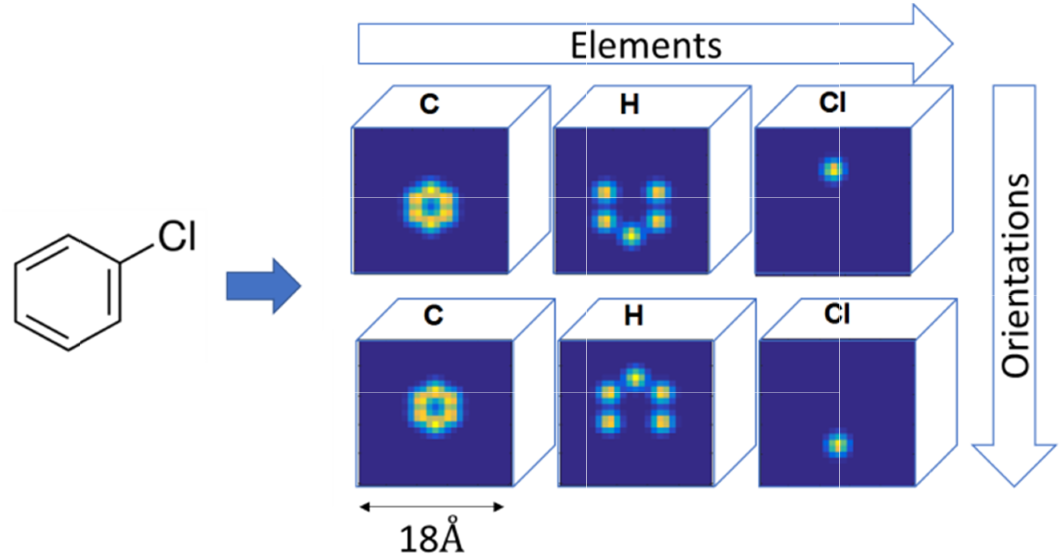
Representing molecules as 5D inputs. Molecules are converted into 3D grids by having each atom as the center of a 3D Gaussian distribution. Just as 2D images have RBG channels, our 3D molecules have one channel for each element. Different orientations of the molecules made up the fifth dimension. In this example, since the molecules are converted into a grid of size 18 Å, with the resolution of 1 Å, with three elements and two orientations, the input would be of size 18^3^·3·2.

### DeepNose autoencoder can be trained to accurately represent the space of molecules

To take advantage of the large dataset containing ~10^7^ unlabeled molecular structures (PubChem3D^19^), and to produce the compact representation of the space of molecules, we designed a stacked autoencoder^20,21^ to accurately represent molecular structures. Autoencoder consists of two convolutional neural networks: an encoder, which converts each molecular structure into a feature vector and a decoder to recover the structure (Figure 2A). We restricted the feature vector to have a lower dimensionality than the input, and therefore induced the network to learn a latent representation that captures the most statistically salient information in the molecular structures. We refer to this latent representation as ‘DeepNose features’. Because it is not necessary and unfeasible to present all available molecular shapes to the Autoencoder, we restricted the training set to 10^5^ molecules randomly selected from PubChem database. We verified that the result does not depend on the selection of molecules. After training using the set of 10^5^ molecules and about 4 · 10^4^ iterations, we achieved a testing set Pearson correlation of r = 0.98 (Figure 2B). Figure 1C shows the network performance of the testing set, calculated at various points during training. Even though the latent representation used ~ 45 times fewer neurons than the input, it can capture the majority of the input information and generalize to novel molecules. We also calculated the correlation coefficients specific to each molecular channel. We observed that C and H channels learn substantially faster than the other channels. This might be due to the higher frequency of C and H atoms in the training set. Nonetheless, autoencoder was able to reconstruct atoms of every element to a correlation of r > 0.95 (Figure 2B-D). Overall, our autoencoder network was capable of accurately replicating small molecules in both our training and testing sets based on the restricted number of features.

**Figure 2.**
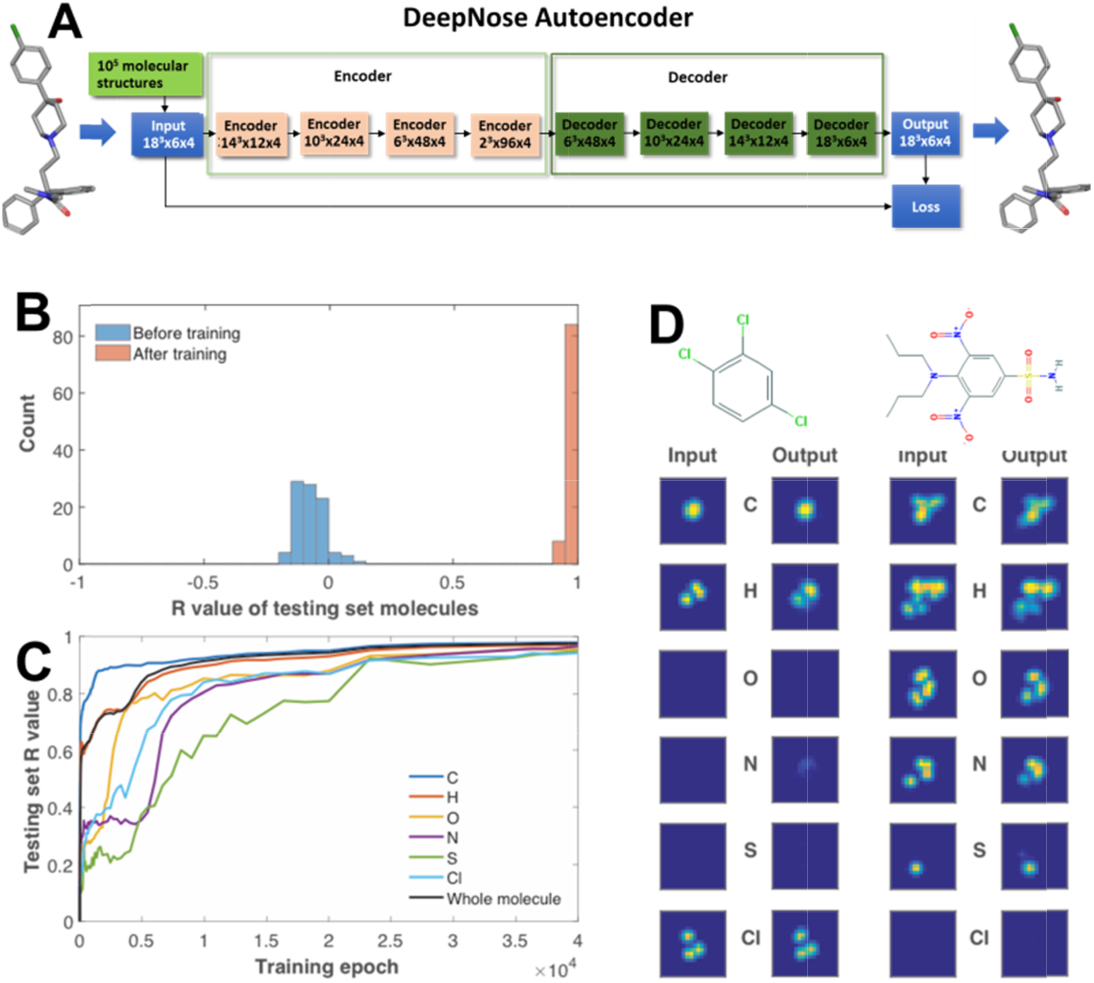
The architecture and performance of DeepNose autoencoder. (**A**) The autoencoder consists of an encoder and a decoder. The encoder transforms the input into a latent representation, while the decoder performs the reverse operation to reconstruct the input molecule. The autoencoder is trained to minimize the difference between the input and the reconstruction. (**B**) Histogram of the correlation coefficient between the original and reconstructed testing molecules before (blue) and after (red) training for 4·10^4^ epochs. (**C**) At different points during training, we calculated the mean correlation coefficient of the molecules in the testing set (black). We also calculated the correlation coefficient separately for individual element: carbon (blue), hydrogen (red), oxygen (yellow), nitrogen (purple), sulfur (green) and chloride (cyan). (**D**) Two examples of molecules reconstructed by the autoencoder are shown: trichlorobenzene (left) and oryzalin (right). Each box is a 2D projection of the 3D grid of individual elements.

### DeepNose features accurately predict water solubility

Are DeepNose features relevant to the physico-chemical properties displayed by molecules? To test the features extracted by autoencoder, we used ESOL dataset^22^ that contains information about water solubility for 1144 molecules to train a neural network classifier. The classifier is composed of the convolutional encoder layers from autoencoder, the consolidation layer that averages responses of 96 features of autoencoder over remaining spatial and orientation variables, and three fully connected classifier layers (Figure 3A). The goal of the consolidation layer is to represent the output of olfactory sensory neurons. In mammalian olfactory system, each olfactory sensory neuron produces only one type of OR and conveys information about the odorant to the olfactory networks^2^. To implement this constraint in our network, since autoencoder features are purported to represent activities of an individual *in silico* OR type, and since responses of olfactory sensory neurons are not expected to be sensitive to the presentation angle and position of each ligand molecule we averaged autoencoder feature activity over ligand spatial and orientation variables. To represent each molecule as input we used its 3D structure replicated at 320 orientations that uniformly sampled three solid rotation angles. Each molecule was therefore defined as a 18^3^ x 6 x 320 5D structure. The classifier receives each molecule’s 5D input structure, generates the set of molecular features using the encoder layers of DeepNose autoencoder, averages these features over space and orientation and passes responses of 96 olfactory ‘sensory neuron’ classes to the fully connected classifier layers.

**Figure 3.**
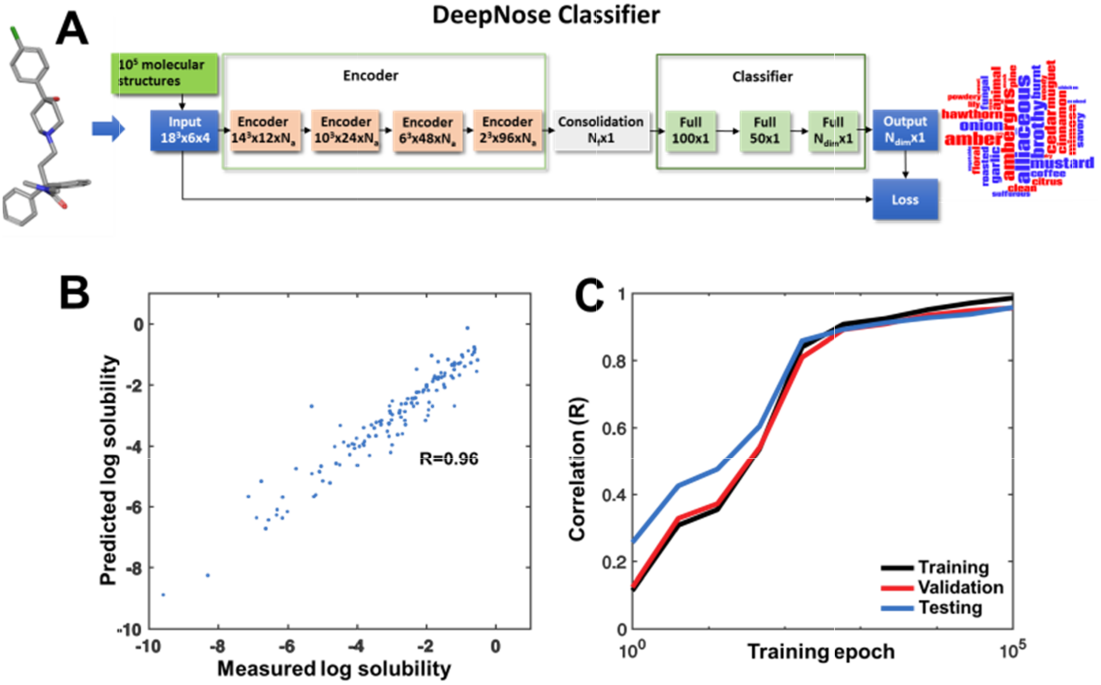
The architecture and solubility prediction performance of DeepNose classifier. (**A**) The classifier consists of three encoder layers, one consolidation layer and three fully-connected layers. The three encoder layers copied the weights from the trained autoencoder’s first three layers; the consolidation layer integrate DeepNose features of N_a_ number of orientations; and the fully-connected layers transformed the features into the bioactivity of N_f_ dimensions. The N_f_ values of the ESOL, Good Scents and Flavornet datasets are 1, 25 and 10, respectively. (**B**) Mean correlation coefficient of training (black), validation (red) and testing (blue) molecules for the ESOL data set at different points during training. (**C**) Scatter plot showing the measured versus predicted log solubility of the testing set after training for 10^5^ epochs (R=0.96).

We trained the classifier using 10^5^ iterations on the training set. We evaluated the performance of the testing set at various time points; the results are shown in Figure 3B and C. DeepNose features, when paired with a neural networks classifier, are predictive of the water solubility with the accuracy of R « 0.96.

### Analysis of olfactory perceptual spaces

It was previously argued that human olfactory space can be viewed as continuous, curved and low-dimensional manifold^23^. We used two olfactory perceptual datasets, Flavornet^24^ and Good Scents^25^ to embed the space of olfactory perceptual objects. These two datasets contain information about human responses about large number of molecules (738 and 3826 respectively) that are represented by a number of semantic descriptors (197 and 580). Because these responses are binary and very sparse, for the purposes of obtaining a reliable predictor, it is more relevant to obtain a low-dimensional approximation of the dataset and to represent each molecule by a dense coordinate along each of the dimensions. To reduce the dimensionality of each of the datasets, we used the Isomap algorithm^26^. Semantic descriptors that are significantly enriched in the positive or negative directions of the first three dimensions are illustrated by word clouds in Figures 4C-E and 5C-E. The first perceptual dimension with the highest included variance has been previously argued to represent molecules’ pleasantness^27^. If were the case, we expect to see a strong correlation of the first perceptual coordinates between molecules that are shared between these two datasets. Indeed, the first perceptual dimensions of the overlapping set of molecules from Flavornet and Good Scents dataset have the highest correlation coefficient amongst all dimension pairs (R = 0.54). Overall, we reduced dimensionality of two discrete semantic datasets of molecules describing human olfactory percepts evoked by these molecules and found that the principal dimensions in both datasets are consistent for overlapping molecules.

**Figure 4.**
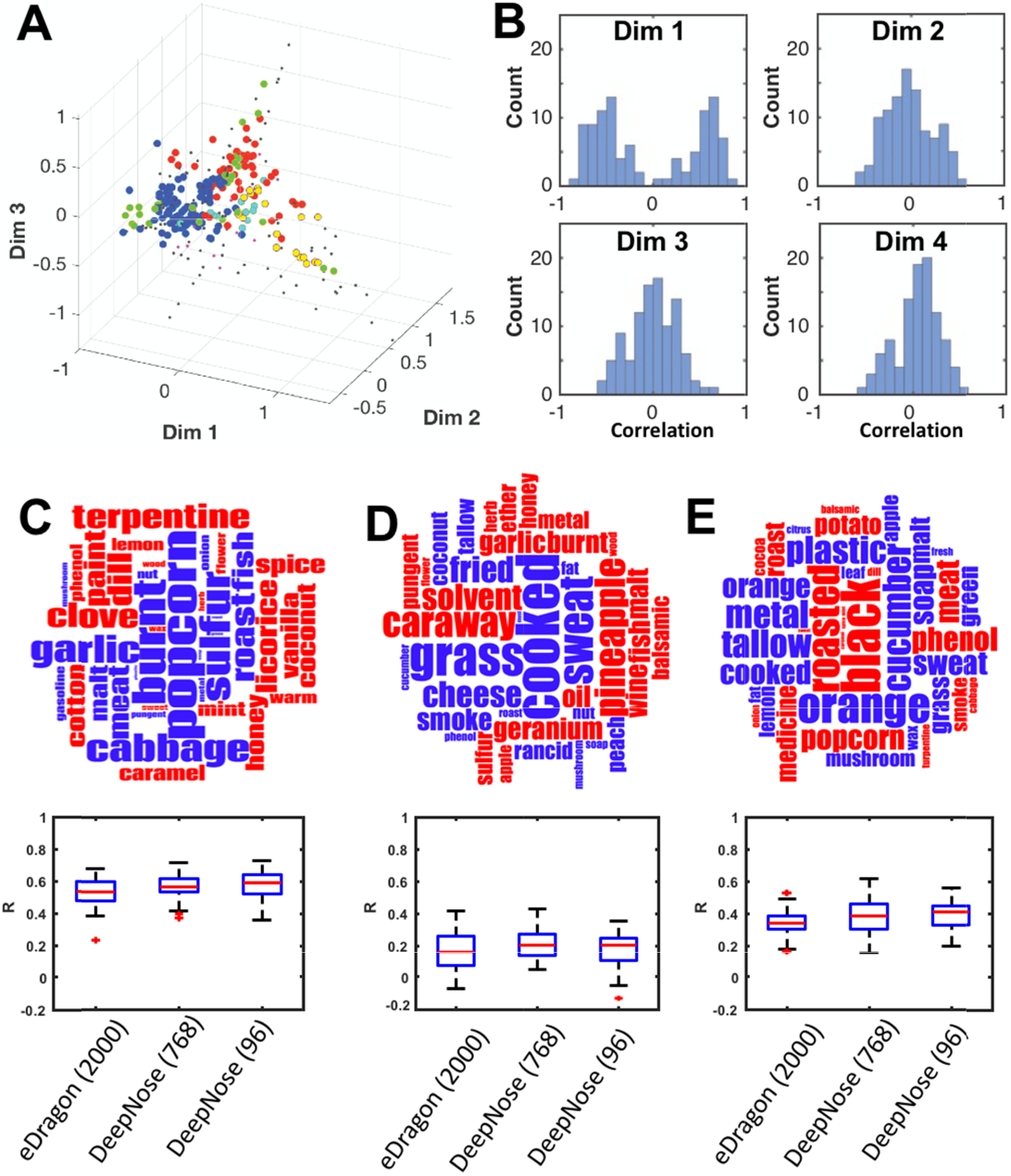
Construction and prediction of the perceptual dimensions from the Good Scents dataset. (**A**) Position of odorant molecules in the first three Isomap perceptual dimensions. (**B**) Correlation coefficient of the Isomap dimensions obtained from two halves of the dataset. (**C-E**) Word clouds showing words significantly enriched in the positive (blue) or negative (red) direction of the first, second and third Isomap dimensions. Prediction accuracy of the first, second and third Isomap coordinates using DeepNose features or E-Dragon chemical descriptors (bottom).

We then tested how robust each dataset’s ISOMAP dimensions are to the selection of the set of molecules (resampling). To this end, we selected two random subsets from each dataset with a small overlap (100 molecules), performed low dimensional embedding for each subset separately and compared Isomap dimensions for the common set of molecules. A large correlation between two subsets (R~ ± 1), would imply that the given Isomap dimension is robust to the selection of molecules. For Good Scents dataset, we found that the first three dimensions are more consistent compared to other dimensions (Figure 5B) suggesting that these dimensions are robust to dataset resampling. For Flavornet, we found that only the first dimension is consistent between the two subsets, and that other dimensions appear uncorrelated between subsampled datasets (Figure 4B). Overall, we suggest that the first dimension in both datasets is both consistent between them and is robust to the selection of molecules. Good Scents contains more robust to resampling dimensions (3 vs 1) which can be explained by more molecules included in Good Scents compared to Flavornet (3826 vs. 738). Interestingly, we also found that the second dimension of Good Scents database is correlated with the third dimension of Flavornet (R~0.4). Each of the three principal Isomap dimensions is represented by the word clouds in Figure 4 and 5.

**Figure 5.**
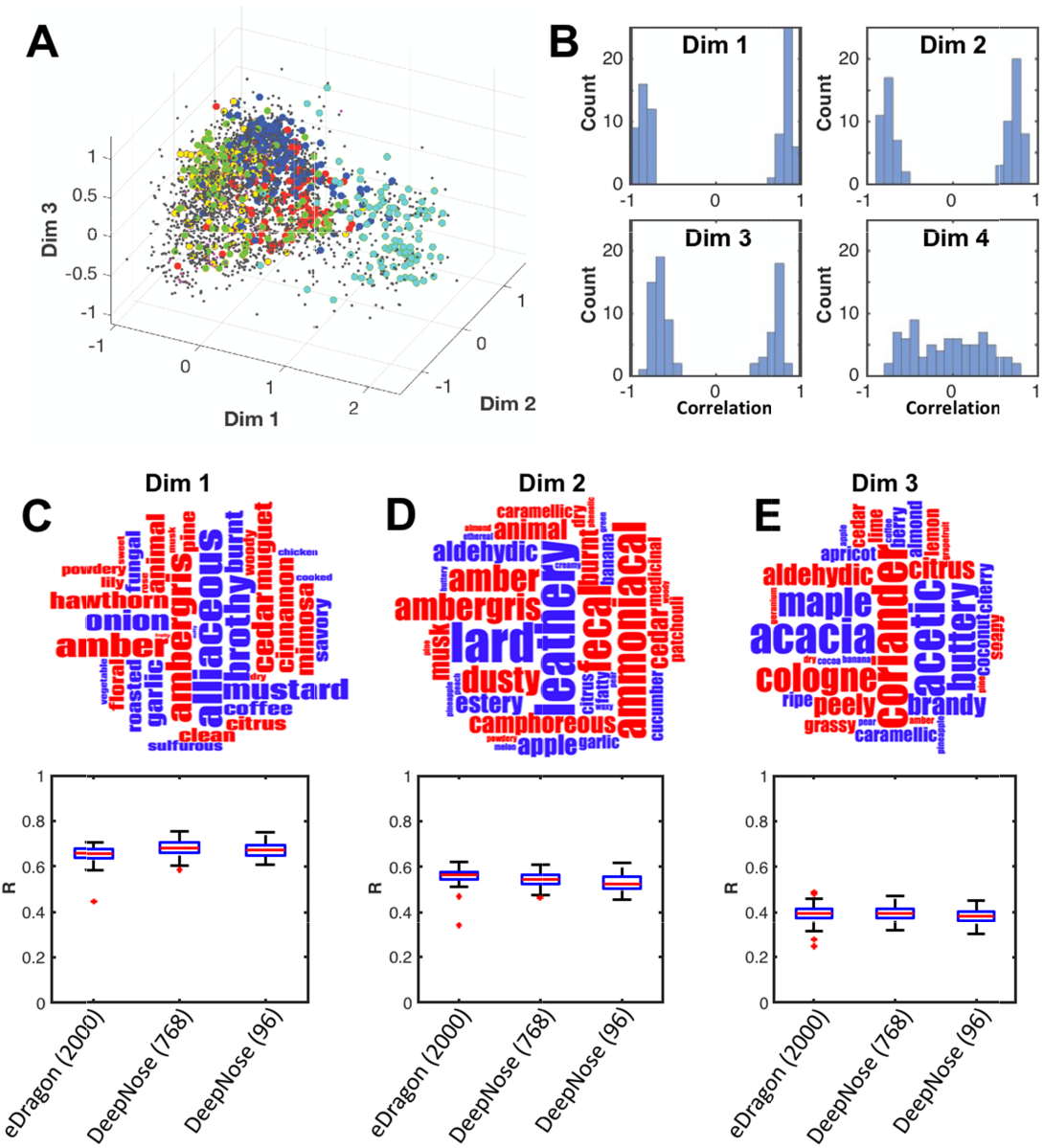
Construction and prediction of the perceptual dimensions from the Flavornet dataset. (**A**) Position of odorant molecules in the first three Isomap perceptual. (**B**) Correlation coefficient of the Isomap dimensions obtained from two halves of the dataset. (**C-E**) Word cloud showing words significantly enriched in the positive (blue) or negative (red) direction of the first, second and third Isomap dimensions. Prediction accuracy of the first, second and third Isomap coordinates using DeepNose features or E-Dragon chemical descriptors (bottom).

### DeepNose features predict perceptual dimensions with comparable accuracy to E-Dragon descriptors

We then used our classifier network to predict human percepts (Isomap dimensions) based on 3D molecular shapes. For each molecule in the Flavornet and Good Scents datasets, we used the trained encoder layers and the consolidation layer to produce a feature vector. These feature vectors were then used as inputs to a fully connected feed-forward neural networks (Figure 3A). To benchmark DeepNose features, we used E-Dragon chemical descriptors often used in olfactory predictions^16,18^, for comparison. E-Dragon features include a diverse set of ~5000 molecular descriptors ranging from simple, such as molecular weight and atomic composition to more complex, such as topological molecular indexes^18^, that summarize the variables developed in computational chemistry to characterize molecules. We included approximately 2000 E-Dragon descriptors that had non-trivial values for the molecules used. In case of E-Dragon features, they were computed for each molecule and used as inputs to the feedforward network similar to the classifier layer in Figure 3A. We find that DeepNose features, when paired with classifier networks, gave predictions of accuracy comparable to E-Dragon descriptors. For example, for the first dimension in Good Scents database (Figure 5C), the median correlation between predicted and observed values is between R=0.67-0.68 if DeepNose features are used versus R=0.66 for E-Dragon features. For the first dimension of Flavornet, the correlations are between R=0.57-0.59 for DeepNose features compared to R=0.54 for E-Dragon. Overall, we suggest that DeepNose autoencoder is able to automatically discover useful chemical features from 3D shapes of molecules alone. These features are useful in predicting human olfactory percepts.

## Discussion

In this study, we present the first description of applying neural networks to directly encode 3D information about molecular conformations. Our main hypothesis is that ORs act as 3D filters that are activated by certain features of 3D molecular confirmations. As such, these features can be extracted and ORs ‘trained’ using conventional machine learning techniques, such as backpropagation. First, we used autoencoder networks to capture the latent space sampled by molecules. Autoencoder, in our implementation, is a convolutional feedforward multilayer neural net which receives a 3D molecular confirmations as inputs and is trained to faithfully replicate these confirmations as outputs. The information about molecular shapes passes through the middle latent layer, which contains about 45 times fewer variables than inputs and outputs. Autoencoder therefore accomplishes its goal using a substantial compression of information. Our assumption is that the ensemble of ORs is under evolutionary pressure to provide a compressed and accurate representation of molecular shapes that is further interpreted by olfactory networks. Under this assumption, we can associate the latent variables produced by our autoencoder with the responses of the ensemble of ORs. If this assumption is true, we can suggest than the process of learning of latent representations mimics the process of evolution of olfactory receptor ensemble. We thus propose a framework within which the evolution of ORs can be viewed as neural network training.

Although DeepNose latent features can be interpreted as OR responses, it is much harder to give a biological interpretation to other layers in our networks. We could argue, for example that the convolutional filters in the first layer of the autoencoder network represent individual amino acids within OR binding pocket, while subsequent layers correspond to assemblies of amino acids of increasing complexity. To make our networks more similar to the biological system, in our classifier network (Figure 3A), we placed a special layer called consolidation. The role of this layer is to average latent variables over positional and orientational degrees of freedom and to compute the output of encoder in a position and orientation independent manner. This layer is therefore intended to replicate the outputs of olfactory receptor (sensory) neurons which are not expected to be sensitive to the ligand position and orientation in space.

We further used the features generated by our network to replicate human olfactory percepts. Because of the need for large number of molecules in the training set, we used large olfactory perceptual datasets (Good Scents and Flavornet) that place molecules into discrete semantic space and contain thousands of molecules. We described olfactory percepts by the salient dimensions of perceptual databases captured by Isomap (Figures 4 and 5). We added several fully connected classifier layers that were expected to give a crude approximation to olfactory networks that have the recurrent structure. We found that the autoencoder’s 96 latent features yield a good predictive value to the first Isomap perceptual dimension (median R~0.67-0.68 for the largest olfactory dataset). These values of correlation were comparable to the one generated by conventional computational chemistry features (E-Dragon, median R=0.66). This finding suggests that DeepNose features represent variables relevant to human olfaction. DeepNose features are computed for each molecules *de novo*, based on this molecule’s 3D shape only. As such, they can be interpreted as the outputs of OR ensemble. It is hard to assume that the olfactory system computes something similar to E-Dragon descriptors. We therefore suggest that our networks provide powerful set of molecular descriptors that can also be interpreted as outputs of biophysical ORs. When information about responses of real OR becomes available for substantial number of ligands, DeepNose features can be adapted to closer resemble their biological counterparts.

## Methods

### Datasets used: PubChem

PubChem is a publicly available dataset that contains more than 200 million unique small molecules^28^. PubChem3D contains pre¬computed 3D conformations of molecules. PubChem3D included up to ten conformations per molecule while attempting to maintain coverage and match experimentally-determined 3D structures^19^. To consider molecules that are odorant-like, we further filtered for those that weigh less than 350 g/mol and contain up to six elements - C, H, O, N, S, and Cl, or "CHONsCi". We randomly selected 10^5^ of these eligible molecules to serve as the training set of the DeepNose autoencoder.

### ESOL

The ESOL dataset contains 1144 molecules with measured solubility^22^. We used this data set to test the ability of DeepNose features to predict water solubility.

### Flavornet

The Flavornet dataset contains 738 odorant molecules associated with one or more perceptual descriptors, such as “flower”, “lemon”, “fish”, etc. There are 197 of these descriptors^24^. We used this data set to test the ability of DeepNose features to predict odorant percepts.

### Good Scents

The Good Scents dataset contains 3826 perfumes associated with one or more semantic perceptual descriptors, such as “fruity”, “woody”, “citrus”, etc. There are 580 of these descriptors. We used this data set to test the ability of DeepNose features to predict odorant percepts.

**Table 1.**
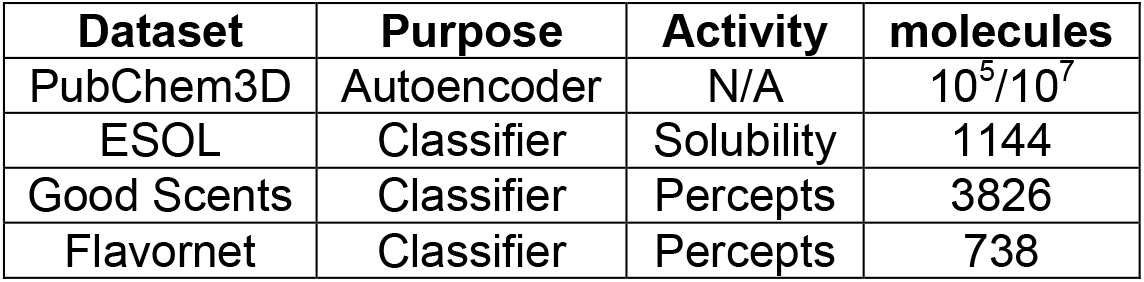
Datasets used.

### Dimensionality reduction of perceptual datasets

We used Isomap^26,27^ to reduce dimensionality of perceptual descriptors in Flavornet and Good Scents datasets. For each data set, we constructed a graph in which each molecule is a node and two molecules are connected by an edge if they share a semantic descriptor. The distance between two connected molecules is calculated based on the overlap between those two molecules’ percepts, according to the following equations:

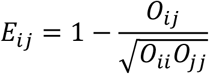

Here *O_ij_* is the number of semantic descriptors shared by molecules *i* and *j*. We then used Isomap to compute geodesic distance between all pairs of molecules and embed them into a low dimensional space. We tested the ability of DeepNose features to predict these perceptual dimensions against molecular descriptors produced by the Dragon 6 software^18^.

One way to visualize what is represented by each Isomap dimension is to identify perceptual descriptors that are enriched in the positive or negative directions of particular dimensions. For example, if a descriptor is associated with N molecules, that descriptor is mapped to N values on the first Isomap dimension. We compared the distribution of these N Isomap values associated with each semantic descriptor to the distribution of all descriptors using the non-parametric Mann-Whitney U test. We obtained a set of semantic descriptors that are significantly enriched in the positive or negative directions of each Isomap dimension. These descriptors are displayed as word clouds.

Second, we tested how robust Isomap dimensions are to subsampling within each dataset. To do that, we split the Good Scents or Flavornet in half and performed the above procedure to approximate the perceptual space in each set. We repeated this analysis for 100 times and looked at the correlation between the two subsets for each dimension. For each dimension, if correlations are near 1 or −1, this Isomap dimension may be considered robust to molecular composition. The distributions of these correlations are shown in Figures 4B and 5B.

Since 520 molecules are present in both Flavornet and the Good Scents datasets, we also tested how much these two datasets correlate for these molecules. We calculated the correlation coefficient between Isomap dimensions in two datasets for overlapping set of molecules. We found that Isomap dimensions 1 and 2 of Good Scents is significantly correlated (R=0.4-0.5) with dimensions 1 and 3 of Flavornet, respectively.

### Input representation

Just as images are represented as 2D objects grouped into separate RGB channels, in our approach, molecules are presented to the network as 3D objects grouped into six channels corresponding to chemical elements (C,O, N, S, Cl, or ‘CHONSCl’). Spatial distribution of atoms of the same type is represented by a 3D grid. We used grids that are 18 Å on each side, with a resolution of 1 Å. Each atom is a 3D Gaussian distribution centering around its xyz-coordinates provided by PubChem3D^19^. We use a fixed standard deviation of 1 Å for all elements. To incorporate different orientations of the molecules, we sample one rotation angle for every training instance of the molecules. The summary of our input representation is summarized in Figure 1.

### Training DeepNose autoencoder

The goal of autoencoder is to produce a compact representation of the space of molecules. It consists of two convolutional neural networks: an encoder, which converts each molecular structure into a feature vector and a decoder to recover the structure. We restricted the feature vector to have a lower dimensionality than the input, and therefore induced the network to learn a latent representation that captures the most statistically salient information in the molecule’s structures. We refer to this latent representation as ‘DeepNose features’.

The network architecture is presented in Figure 2A. One training instance can include one or more orientations, and the same weights are used for each orientation. Starting from random weights, the autoencoder is trained to minimize the difference between the original structure and its reconstruction using backpropagation algorithm^15^. We trained the network using 4· 10^4^ batches, using a batch of 48 molecules for each iteration. We used a training set of randomly selected 10^5^ PubChem3D^19^ molecules and cross-validated our network with a testing set of randomly selected 92 molecules in the ESOL dataset. We evaluated the network performance by calculating the pixel-to-pixel error and correlation between the original molecules and the reconstructed molecules in the testing set.

### DeepNose classifier

The input of the classifier is a 4D object, similar to that of the autoencoder. We pair the trained encoder layers of our DeepNose autoencoder with fully-connected feed-forward layers to predict bioactivities of molecules, including solubility and perceptual coordinates. Since molecules adopt multiple conformations and orientations, we made a simplifying assumption that the different orientations contribute with equal probability to its bioactivity. We therefore add a “consolidation layer” which averages network activity over multiple orientations of the same molecules within the same latent feature.

Classifier network architecture is summarized in Figure 3A. Starting with random weights, the classifier is trained to minimize the difference between the target bioactivity and the network output using backpropagation. We split the dataset into three groups, 70%, 15%, and 15% for training, validation and testing, respectively. We evaluate the model by calculating the error and correlation between actual and predicted bioactivities of molecules in the testing set.

